# Metabolomics and transcriptomics reveal the response mechanisms of *Mikania micrantha* to *Puccinia spegazzinii* infection

**DOI:** 10.1101/2022.06.10.495692

**Authors:** Xinghai Ren, Guangzhong Zhang, Yujuan Gu, Mengjiao Jin, Michael D. Day, Huai Liu, Wanqiang Qian, Bo Liu

## Abstract

*Mikania micrantha* is one of the 100 worst invasive species globally and can cause significant negative impacts to agricultural and forestry economics, particularly in Asia and the Pacific region. The rust *Puccinia spegazzinii* has been used successfully as a biological control agent in several countries to help manage *M. micrantha*. However, the response mechanisms of *M. micrantha* to *P. spegazzinii* infection have never been studied. To investigate the response of *M. micrantha* to infection by *P. spegazzinii*, an integrated analysis of metabolomics and transcriptomics was performed. The levels of 101 metabolites, including organic acids, amino acids and secondary metabolites in *M. micrantha* infected with *P. spegazzinii*, were significantly different compared to those in plants that were not infected. The relative amounts of L-leucine, L-ioleucine, L-tryptophan, histidine, L-phenylalanine and L-citrulline were increased under *P. spegazzinii* infection. These may convert to intermediates within the TCA cycle, participating in energy biosynthesis. In addition, phytoalexins, such as maackiain, nobiletin, vasicin, arachidonic acid and JA-Ile, were synthesized and accumulated in *M. micrantha* in response to *P. spegazzinii* infection. A total of 4,978 differentially expressed genes were identified in *M. micrantha* infected by *P. spegazzinii*. Many key genes of the Mitogen-activated protein kinase (MAPK) signalling pathway, showed significantly higher expression under *P. spegazzinii* infection. These results not only help us to understand the response of metabolites and gene expression in *M. micrantha* following infection by *P. spegazzinii*, but also help us understand the potential for the continuous biological control of *M. micrantha* using *P. spegazzinii*.

## Introduction

*Mikania micrantha* Kunth (Asteraceae), commonly called ‘mile-a-minute’, is a rapidly growing vine native to tropical America (Day *et al*., 2016; CABI, 2019). It is considered one of the world’s worst weeds (Love *et al*., 2014), invading many countries in Asia and Oceania (Day *et al*., 2016; CABI, 2019;) and has been listed in China as a plant species with national quarantine concerns by the Forestry Administration (Zhang *et al*., 2004; Zhang *et al*., 2019). *Mikania micrantha* has been reported in seven provinces in southern China, with the main infestations in Guangdong, Guangxi and Yunnan provinces (Shen *et al*., 2018), and affects biodiversity and a wide range of agriculture and forestry enterprises, reducing productivity through competition (Day *et al*., 2016). *Mikania micrantha* is also known as a plant killer because its rapid growth enables it to completely smother crops or trees and block sunlight preventing flowering and fruiting (Zhang *et al*., 2004; Day *et al*., 2016).

The rust fungus, *Puccinia spegazzinii* de Toni (Uredinales: Pucciniaceae) has been introduced into several countries, including China to help control *M. micrantha* (Day *et al*., 2016). *Puccinia spegazzinii* is a microcyclic and autoecious rust, with a life cycle of 19-21 days (Ryan & Ellison, 2003). Teliospores and basidiospores of *P. spegazzinii* have only been recorded in the field (Ellison *et al*., 2008). *Puccinia spegazzinii* is known to attack only *M. micrantha* and to a much lesser extent, *Mikania cordata* (Burm. F.) B. L. Robinson. Its potential as a biological control agent is because it is particularly damaging to the leaves, stems and petioles of *M. micrantha*, reducing growth rates and flowering and can tolerate a wide range of environmental conditions in which *M. micrantha* grows (Barreto & Evans, 1995; Day *et al*., 2013a, b; Day & Riding, 2019).

Rust fungi (Pucciniales) are one of the largest groups of plant pathogens and the most damaging to plants in the world (Kemen *et al*., 2015; Lorrain *et al*., 2019). Infected plants can produce a range of defence responses to a pathogen (Feng *et al*., 2015). The amount of adenosine triphosphate (ATP), which plays an important role in metabolite biosynthesis, signal transduction, and material transport can be increased. These secondary metabolites play vital roles in performing or having passive physical and chemical barriers against pathogens. For instance, polyamines could change the mesophyll cell pre-penetration and penetration resistance mechanisms, e.g. as seen in oats in response to infection by *Puccinia coronata* f. sp. *avenae* W.P. Fraser & Ledingham (Pucciniaceae) (Montilla-Bascon *et al*., 2016).

Other responses relate to the phenylpropanoid, flavonoid and isoflavonoid metabolic pathway genes, which are involved in the production of phytoalexins as seen in *Medicago truncatula* Gaertn (Fabaceae) following infection by *Phakopsora pachyrhizi* Syd. & P. Syd. (Phakopsoraceae) (Ishiga *et al*., 2015). In addition, the accumulation of phenolic, rutin, glucosinolate, flavonoid, fatty acid and alkaloid phytoalexins in plants could promote host defence against pathogens (Maddox *et al*., 2010; Amil-Ruiz *et al*., 2011; Joosten & van Veen, 2011). Defence-signalling networks, such ethylene and the salicylic acid pathways, are also triggered when pathogens attempt to infect host plants (Yang *et al*., 2014) and several gene families, such as the MCM1, WRKY, MYB, and bZIP families, are involved in the response of plants to pathogens (Al-Attala *et al*., 2014; Wang *et al*., 2017; Zhu *et al*., 2018; Rathod *et al*., 2020).

In this study, we have characterized *M. micrantha* as a model host pathosystem for *P. spegazzinii* to investigate the molecular mechanisms of host response to infection. Using transcriptome and metabolome analyses, we show how levels of essential amino acids, ATP and phytoalexins in *M. micrantha* change in response to infection by *P. spegazzinii*. In addition, we aim to show how genes related to biotic stress in *M. micrantha* were expressed following *P. spegazzinii* infection. This work will help understand the potential for the continuous use of *P. spegazzinii* in assisting in the management of *M. micrantha* in many countries.

## Materials and Methods

### Rust inoculation and sample collection

*Puccinia spegazzinii* IMI 393075 was imported into China from Australia in December 2019 and a culture was maintained on potted *M. micrantha* plants at the Agricultural Genomics Institute at Shenzhen, Chinese Academy of Agricultural Sciences. *Mikania micrantha* plants were grown in a greenhouse with lighting (day: 16 hours and night: 8 hours) and a daily temperature range of 20-35°C. For inoculation, plant tissue (leaves, stems, petioles) with mature *P. spegazzinii* pustules, indicated by a coppery-brown appearance, was suspended over healthy plants in a sealed plastic chamber maintained at 22°C, 100% humidity, for 48 hours of darkness (for details, see Ellison *et al*., 2008; Day & Riding, 2019). After 48 h, the plants were removed and placed in an insect cage in a greenhouse (25-35°C, relative humidity, ambient lighting) for subsequent pustule development. After 17 days following inoculation, infected leaves with obvious well-developed pustules, were collected in three biological replicates. The pustules were cut from the leaves to eliminate the metabolites of the rust contaminating the leaf samples and discarded. Leaves of uninfected plants were also collected in three biological replicates to serve as controls. The collected leaf samples were placed in liquid nitrogen for quick freezing and then stored at −80°C for further experiments.

### Metabolite response of *M. micrantha* after infection by *P. spegazzinii*

Following freezing by liquid nitrogen, the samples of infected leaves and uninfected leaves were ground into powder. A subsample of 100 mg powder from each of the three samples of infected and uninfected leaves was weighed and the homogenate was resuspended with prechilled 500μL 80% methanol and 0.1% formic acid by vortexing well. The samples were incubated on ice for 5 min and were then centrifuged at 15000 g, at 4°C for 10 min. The supernatant was diluted using LC-MS grade water so the final concentration was 53% methanol. The samples were subsequently transferred to fresh Eppendorf tubes and were centrifuged at 15000 g, at 4°C for 20 min. The supernatant was injected into the Liquid Chromatography–Tandem Mass Spectrometry (LC-MS/MS) system analysis.

Based on novogene Database (novoDB, in-house database), the Multiple Reaction Monitoring (MRM) was used to detect the metabolites in each sample. The product ion (Q3) was used for metabolite quantification. The parent ion (Q1), Q3, retention time (RT), declustering potential (DP) and collision energy (CE) were used for metabolite identification. The data files generated by HPLC-MS/MS were processed using the SCIEX OS Version 1.4 to integrate and correct the peaks. The main parameters were set as follows: minimum peak height, 500; signal/noise ratio, 5; gaussian smooth width, 1. The area of each peak represents the relative content of the corresponding substance.

These metabolites were annotated using the Kyoto Encyclopedia of Genes and Genomes (KEGG) database (http://www.genome.jp/kegg/), Human Metabolome Database (HMDB) database (http://www.hmdb.ca/) and Lipid maps database (http://www.lipidmaps.org/). We applied univariate analysis (t-test) to calculate the statistical significance (P-value). The metabolites with a P-value < 0.05 and a fold change (FC) ≥ 2 or ≤ 0.5 were considered to be differential metabolites.

### Establishment of cDNA library and differentially expressed gene (DEG) analysis

The total RNA of *M. micrantha* leaves infected by *P. spegazzinii* and leaves from uninfected plants was extracted using an RNA extraction kit, and the quality of the extracted RNA was checked using a Nanodrop 2000 spectrophotometer. The integrity of total RNA was checked using formamide denaturing gel electrophoresis. mRNA was isolated from the total RNA using Dynabeads Oligo (dT) 25 isolation beads. The RNA of the extracted sample was used to build a cDNA library by a reverse transcription kit based on the manufacturer’s instruction (NEBN_ext_^®^ Ultra^™^ RNA Library prep Kit for Illumina^®^).

After establishing a cDNA library, the concentration of purified cDNA was preliminarily quantified by a Qubit2.0 fluorometer. The insert size of the cDNA library was checked by an Agilent 2100 bioanalyser. To ensure a quality library, the concentration of purified cDNA was accurately quantified using Q-PCR. The cDNA library was sequenced on the Illumina sequencing platform, using paired-end (PE) technology, in which 150 bp PE reads were generated. The quality control of RNA-Seq reads was assessed by Trinith (Chen *et al*., 2019).

To obtain localization information for the reads in the reference genome, clean reads were compared with the reference genome of *M. micrantha* (GCA_009363875.1) (Liu *et al*., 2020), using HISAT2-2.0.5 (Pertea *et al*., 2016), and the expression level was calculated using the fragments per kilobase million (FPKM) method. The differentially expressed genes (DEGs) were analysed using the DEseq2 package version 3.8.6 (Love *et al*., 2014). Genes with a | log_2_ fold change | > 1 and false discovery rate (FDR) <0.05 were considered DEGs. The KEGG enrichment analysis of functional significance terms based on the KEGG (http://www.kegg.jp/kegg/pathway/html) database was conducted using a hypergeometric test to find significant KEGG terms in DEGs for comparison with the genome background.

### Contents of ATP determination following *P. spegazzinii* infection

Following freezing by liquid nitrogen, the samples of infected leaves and uninfected leaves were ground into powder. A subsample of 100 mg powder from each of the three biological samples of infected and uninfected leaves was weighed and 1ml buffer solution was added and homogenized in an ice bath using an electric tissue grinder. The mixed extracts were then vortexed and centrifuged at 8000 g and 4°C for 10 min. The supernatant was placed into a new Eppendorf tube, and 500 μL trichloromethane was added before being mixed thoroughly, and centrifuged at 10000 g and 4°C for 3 min. The upper suspension of each sample was kept and incubated on ice for subsequent detection of ATP content in infected and uninfected leaves using an ATP content detection kit (Beijing Solarbio Technology Co., Ltd.).

## Results

### Metabolite response of *M. micrantha* after infection by *P. spegazzinii*

A total of 101 significant differentially abundant metabolites (23.88% of total metabolites) were identified in the leaves of *M. micrantha* in this study. Of these, 39 metabolites had significantly higher levels in infected leaves compared with the uninfected *M. micrantha* samples, while 62 metabolites had significantly lower levels following infection by *P. spegazzinii*, compared to the uninfected *M. micrantha* samples (**Figure 1A**). The 101 significant differentially metabolites could be grouped into 24 major classes (**Supplementary Table S1**). Flavonoids were the most abundant metabolites among the significantly differential metabolites, accounting for 13.86% of the total significantly metabolites (**Supplementary Figure S1**).

**Figure 1.**
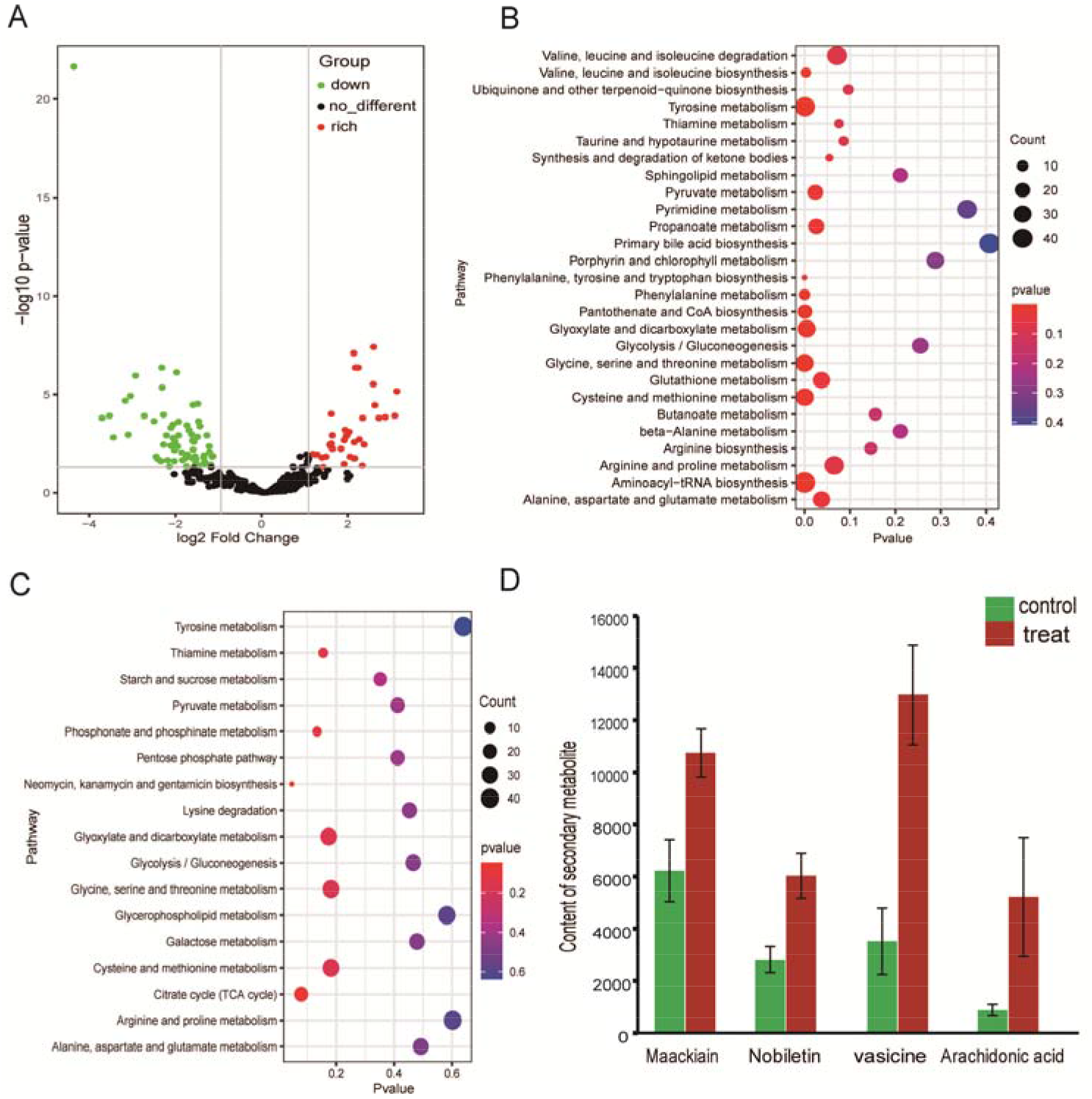
Volcano map and enrichment of differential metabolites. (A) Volcano map of differential metabolites. Red plots indicate the up-regulated metabolites; Green plots indicate the down-regulated metabolites; Black plots indicate No significant difference. (B) KEGG enrichment pathway of up-regulated metabolites. (C) KEGG enrichment pathway of downregulated metabolites. (D) Contents of secondary metabolites of M. micrantha under P. spegazzinii stress. Different letters indicate the level of difference, p<0.05.

Based on the KEGG database, a total of 19 significantly increased metabolites are enriched in essential amino acid synthesis and metabolic pathways, such as phenylalanine, tyrosine and tryptophan biosynthesis; phenylalanine metabolism; valine, leucine and isoleucine degradation; glycine, serine and threonine metabolism (**Figure 1B**). The co-joint KEGG enrichment analysis of the significantly decreased metabolites, showed 17 co-mapped pathways, such as those related to neomycin, kanamycin and gentamicin biosynthesis, citrate cycle (TCA cycle), phosphonate and phosphinate metabolism, thiamine metabolism, and glyoxylate and dicarboxylate metabolism (**Figure 1C**).

The levels of phytoalexin-related metabolites, e.g. maackiain, nobiletin, vasicin and arachidonic acid in *M. micrantha* leaves infected with *P.spegazzinii* were 1.73, 2.14, 3.69 and 5.98 times higher respectively than those in uninfected leaves (**Figure 1D**). The level of the stress-related compound, JA-Ile was 6.40 times higher in infected leaves than that in uninfected leaves (**Figure 1D**).

### Differentially expressed gene (DEG) analysis

Based on the transcriptome analysis between the *P. spegazzinii*-infected group (A) and non-infected (control) group, a total of 4,978 DEGs (15.20% of total genes) were identified, including 2,054 significantly up-regulated genes and 2,924 significantly down-regulated genes (**Figure 2A**). Among these DEGs, a total of 2,078 DEGs are enriched in 133 KEGG pathways (Supplementary Table **S2**). The most significantly enriched pathways of up-regulated DEGs are related to protein processing in endocytosis and plant-pathogen interaction (**Figure 2B**). The expression of genes associated with biosynthesis of terpenoid and flavonoid, such as in terpenoid backbone biosynthesis, isoflavonoid biosynthesis, monoterpenoid biosynthesis, sequiterpenoid and triterpenoid biosynthesis and flavone and flavonol was significantly suppressed (**Figure 2C**).

**Figure 2.**
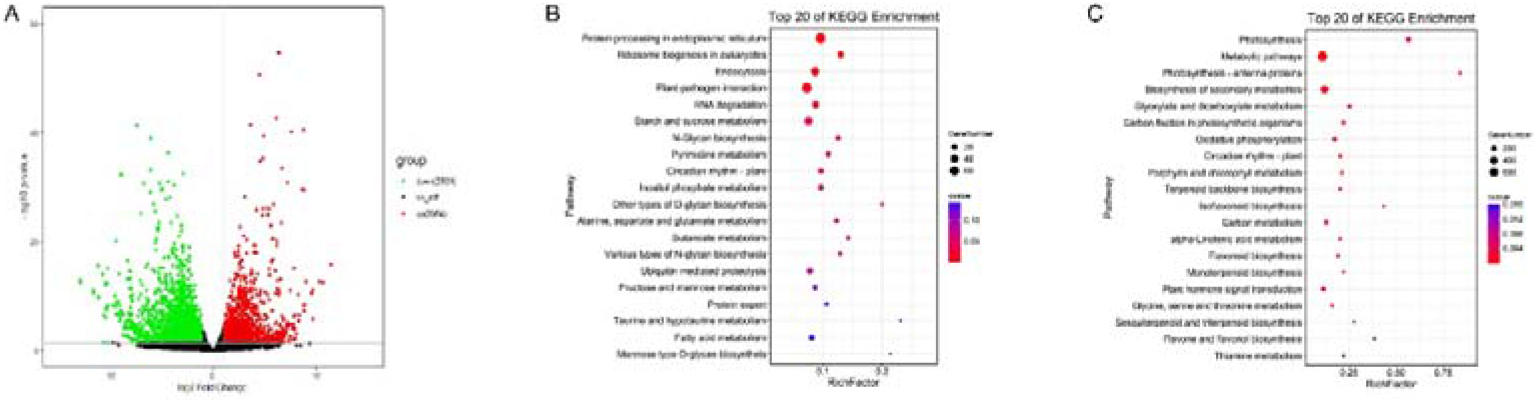
Volcano map and enriched of differential genes. (A) Volcano map of differentially expressed genes; Red plots indicate the up-regulated metabolites; Green plots indicate the down-regulated metabolites; Black plots indicate No significant difference. (B) KEGG pathway enrichment of up-regulated genes; (C) KEGG pathway enrichment of down-regulated genes.

A total of 95 WRKY genes were identified in the *M. micrantha* genome, in which the gene expression of 22 WRKY (23.16% of total WRKY genes) were significantly different following infection by *P. spegazzinii* as determined by the RNA-seq analysis. Among these genes, 12 WRKY genes were significantly up-regulated, including one WRKY1 gene, seven WRKY22 genes, three WRKY33 genes and one WRKY4/5 gene. The gene expression of the three WRKY33 genes were significantly up-regulated by 2.2 times following *P. spegazzinii* infection (**Figure 3** and **Supplementary Table S3**). A total of 6 BAK1 genes were identified in the *M. micrantha* genome, in which the gene expression of 2 BAK1 genes were significantly up-regulated by 1.5 times following *P. spegazzinii* infection (**Figure 3** and **Supplementary Table S3**).

**Figure 3.**
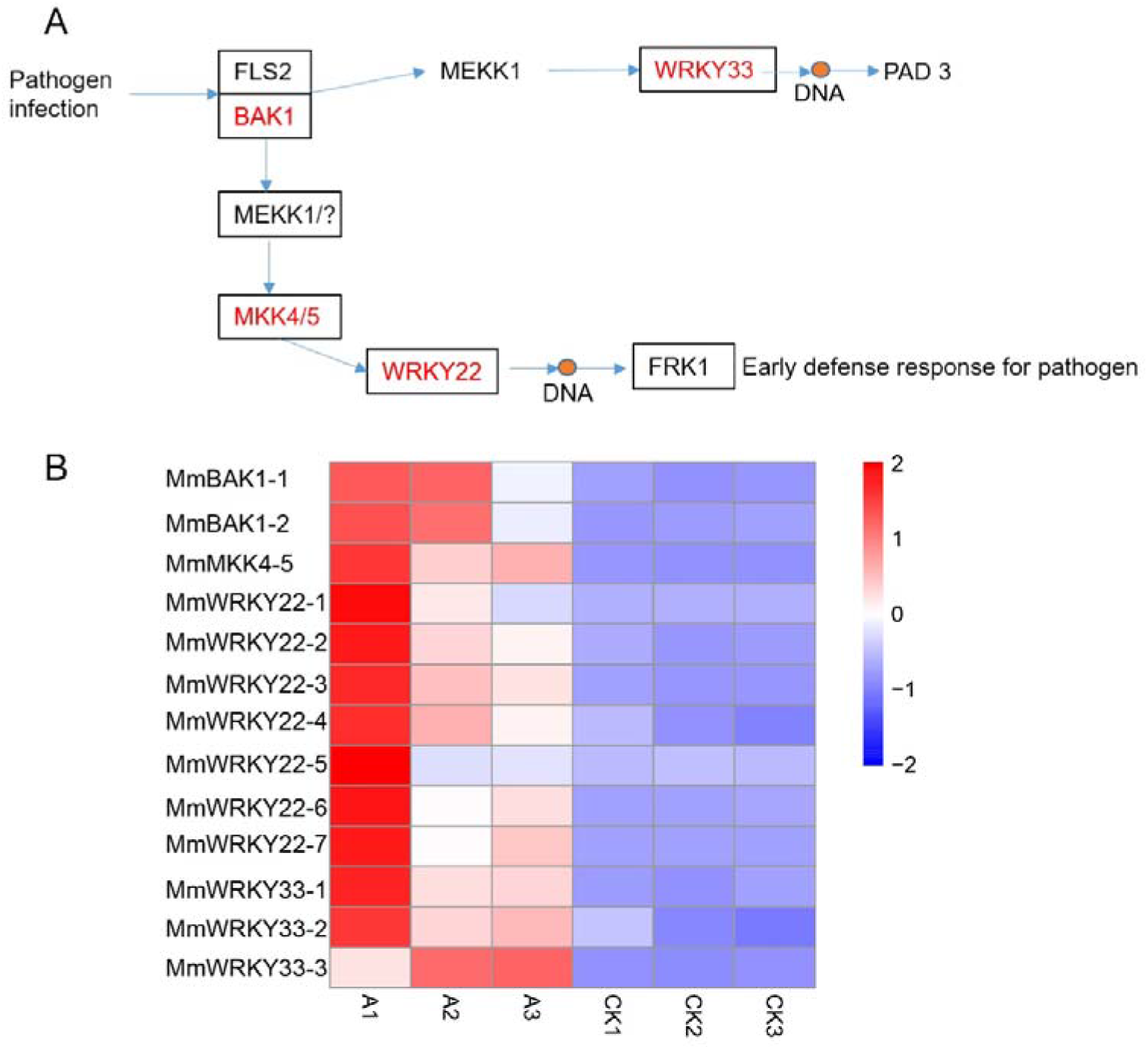
MAPK signaling pathway-plant in *M. micrantha* under infection by *P. spegazzinii*. The gene expression levels were denoted in blue (down-regulated) and red (up-regulated). (A) MAPK signaling pathway-plant; (B) expression pattern of MAPK signalling pathway-plant key genes in *M. micrantha*. The red box indicates upregulated expression.

### Contents of ATP determination following *P. spegazzinii* infection

The gene expression of key amino acids synthesis, such as branched-chain amino acid aminotransferase and 2-isopropylmalate synthase, was significantly up-regulated when *M. micrantha* was infected by *P.spegazzinii* (**Figure 4A** and **4C**). Moreover, the amounts of these amino acids, such as L-leucine, L-isoleucine, L-tryptophan, histidine, L-phenylalanine, and L-citrulline, which can turn into succinyl coenzyme A and acetyl-CoA by multistep transamination and decarboxylation, were also significantly increased after *P.spegazzinii* infection (**Supplementary Figure S2**). Furthermore, the expression of almost all TCA cycle-related genes (6 of 8 key genes) was significantly up-regulated (**Figure 4B,D** and **Supplementary Table S3**), causing the amounts of these amino acids to be significantly increased, promoting the production of ATP following infection by *P.spegazzinii* (**Figure 4E**).

**Figure 4.**
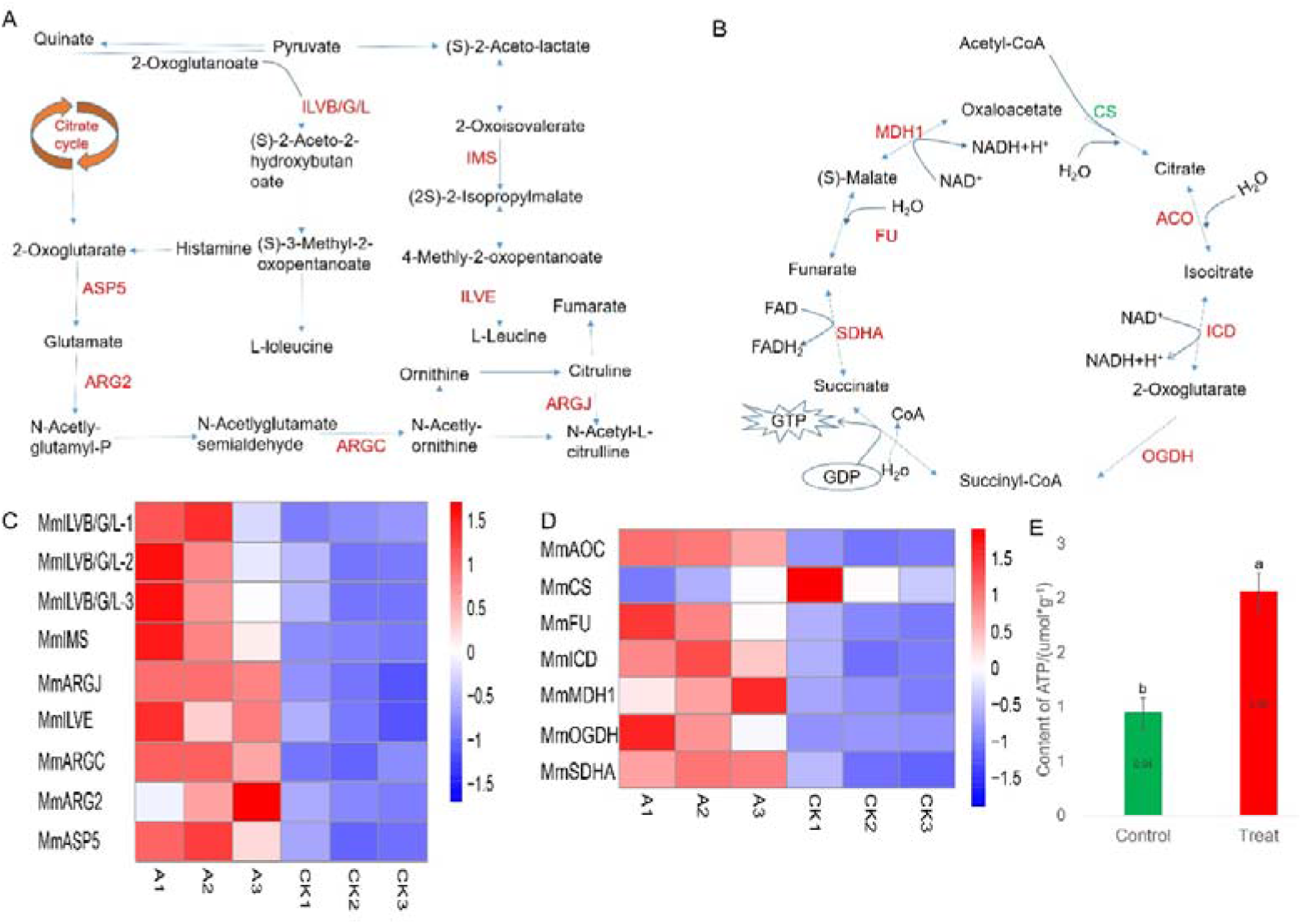
Gene expression of amino acid metabolism and TCA cycle pathway in *M. micrantha* under *P. spegazzinii* stress. (A) Pathway of amino acid metabolism; (B) pathway of the TCA cycle; (C) Expression pattern of key amino pathway genes in M. micrantha; ILVB/G/L: acetolactate synthase I/II/III large subunit; IMS: 2-isopropylmalate synthase; ILVE: branched-chain amino acid; ARGJ: glutamate N-acetyltransferase; ARGC: N-acetyl-gamma-glutamyl-phosphate reductase; ARG2: amino-acid N-acetyltransferase; ASP5: aspartate aminotransferase, chloroplastic. (D) Expression pattern of key TCA cycle genes in M. micrantha; SDHA: succinate dehydrogenase (ubiquinone) flavoprotein subunit; FU: fumarate hydratase, class II; MDH1: malate dehydrogenase 1; CS: citrate synthase; ACO: aconitate hydratase; ICD: isocitrate dehydrogenase; OGDH: 2-oxoglutarate dehydrogenase E1 component. (E) Changes in the contents of ATP by the P. spegazzinii infection; Different letters indicate the level of difference, p<0.05.

## Discussion

There were both positive and negative responses in terms of levels of metabolites and gene expression when *M. micrantha* was infected with *P. spegazzinii*. While most of the metabolites and gene expressions decreased following infection, there were numerous metabolites and genes that had increased, suggesting that infection by *P. spegazzinii* had stimulated compensatory or defence responses. Over one third of the metabolites detected, had significantly higher levels than that seen in uninfected plants. Most of these metabolites pertain to amino acid synthesis and metabolic pathways, suggesting that *M. micrantha* is able to produce more amino acids to maintain growth following infection by *P. spegazzinii*.

The amounts of essential amino acids, ATP and phytoalexins (e.g. nobiletin, maackiain and JA-Ile) were significantly increased following infection by *P. spegazzinii* infection. The stress related compound JA-Ile which plays an important role in response to the biotic or abiotic stressors (Wasternack, 2014), was significantly increased more than 6.4 times following infection by *P. spegazzinii*, again suggesting some sort of defence mechanism.

About 40% of genes identified were up-regulated and most of these were related to protein processing, suggesting the plant may be trying to maintain growth while infected. There were numerous genes e.g. WRKY33 and WRKY22, relating to plant resistance (Jiang *et al*., 2017) that had also increased following infection, suggesting the plant is trying to overcome infection. ATP production also increased following infection, as seen in other studies (e.g. Montilla-Bascon *et al*., 2016).

In addition, the key genes in the TCA cycle and MAPK signalling pathways presented high expression following *P. spegazzinii* infection. These results suggested that when *M. micrantha* was infected by *P. spegazzinii*, it activated a defence response, but this does not necessarily imply resistance.

In previous studies, other plant species have responded to the presence of pathogens by up-regulating metabolites and DEGs (Pitino *et al*., 2017; Tang *et al*., 2018; Wang *et al*., 2018). The innate immune system (Gu *et al*., 2019) and phytoantitoxin (Stevenson & Haware, 1999; Kliebenstein, 2004) of plants are used to respond to pathogen infection through cell permeability, programmed cell death and defence responses (Benaziz, 1967; Savchenko *et al*., 2010; Yao *et al*., 2012; Zlotek & Wojcik, 2014; Zaynab *et al*., 2018). When plants are under biotic and abiotic stress, the WRKY family plays vital roles in stress-induced defensive responses (Jiang *et al*., 2017; Wang *et al*., 2017; Satapathy *et al*., 2018; Wang *et al*., 2019; Rathod *et al*., 2020; Ramos *et al*., 2021).

This study has shown that levels of some genes and metabolites increased following infection by *P. spegazzinii*, suggesting that the plant is trying to compensate or overcome infection. Some of these positive responses relate to resistance which then raises the question of whether *M. micrantha* could ever become resistance to *P. spegazzinii* and therefore, will the rust’s impact as a biological control agent lessen?

Despite these positive changes in levels of metabolites and gene expression, this study, also showed that *P. spegazzinii* can have a negative impact on gene expression, metabolites and metabolic pathways. Most of the metabolites (>60%) and genes (60%) had lower levels following infection, suggesting that infection of *P.spegazzinii* is having some negative impact on *M. micrantha*.

*Puccinia spegazzinii* is currently being used as an effective biological control agent and has been released against *M. micrantha* in many countries (Ellison *et al*., 2008; Winston *et al*., 2014; Day *et al*., 2016; Kumar *et al*., 2018; Day & Riding, 2019). The management of *M. micrantha* using *P.spegazzinii* is seen as a benefit to many land holders, as plants can reshoot after slashing and herbicides are harmful to humans and the environment (Choudhury, 1972; Day *et al*., 2016; Day & Riding, 2019). In both laboratory and field trials, *P. spegazzinii* has been shown to reduce growth rates and flowering of *M. micrantha* (Day *et al*., 2013b). The results in this study at least partially explain what is being observed in the field where growth rates of *M. micrantha* have decreased, offering hope that *P. spegazzinii* will continue to be able to suppress populations of *M. micrantha* in the field.

In this study, we characterized *M. micrantha* as a model host pathosystem for *P. spegazzinii* to investigate the molecular mechanisms of host responses to a pathogen. We have provided the first description genome wide, of the different gene expressions and metabolite levels in the leaves of *M. micrantha* following *P. spegazzinii* infection. Interactions and coevolution between *M. micrantha* and *P. spegazzinii* may influence the overall pathogenicity of rust. Over time, *M. micrantha* may evolve a mechanism to resist *P. spegazzinii*, and the plant could once again become a problem for countries that already have released rust. Alternatively, the rust may develop ways to overcome the plant’s defence responses or resistance. Further studies may provide more insight. The results here certainly provide a better understanding of the interactions between *M. micrantha* and *P. spegazzinii* and the potential of *P. spegazzinii* as a long-term biological control agent for *M. micrantha*.

## Conclusion

In this study, we revealed the response and interaction mechanism of *M. micrantha* to *P. spegazzinii* based on the analysis of plant metabolites and gene expression. The relative levels of essential amino acid, ATP and phytoalexins were synthesized and accumulated in *M. micrantha* following *P. spegazzinii* infection. In addition, many key genes of the MAPK signalling pathway, showed significant high expression under the *P. spegazzinii* infection.

## Data availability statement

Publicly available datasets were analyzed in this study. The RNA-seq data can be found in the NCBI under the accession numbers SRR17139378–SRR17139383.

## Acknowledgements

This work was carried out with the support of projects subsidized by special funds for the National key research and development program of China (2021YFC2600100), science technology innovation and industrial development of Shenzhen Dapeng New District (Grant No. KJYF202001-03).

## Author contributions

X.H.R. and Y.J.G. carried out the sample collection and the inoculation of rust. X.H.R. and G.Z.Z provided technical support for the bioinformatics analysis. M.J.J. detected the contents of ATP in plant leaves. X.H.R. and B.L. wrote the manuscript. M.D.D. revised the paper and provided context in terms of biological control. B.L. W.Q.Q.and H.L. designed the experiments and coordinated the project.

---

**Supplementary Figure S1.**
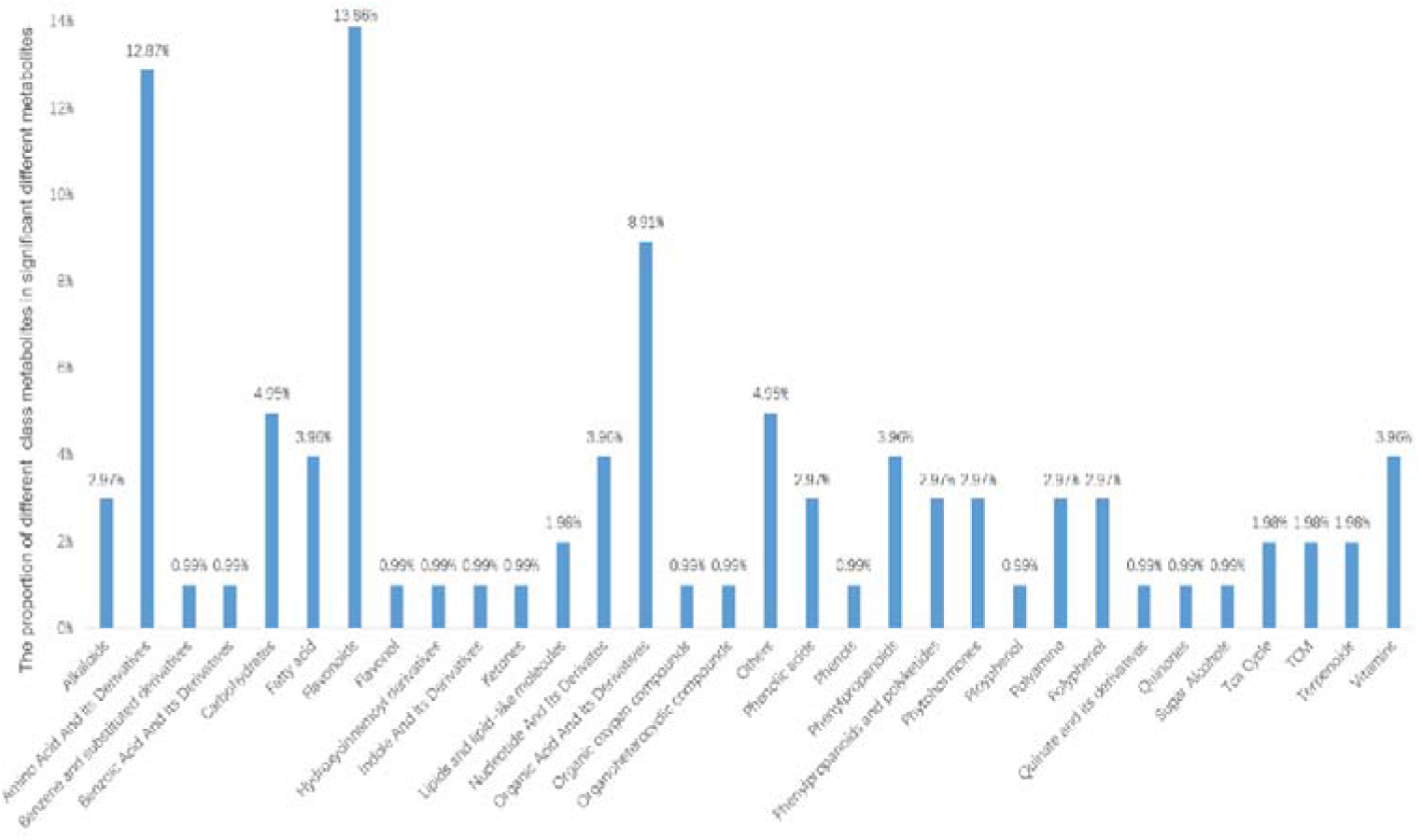
The proportion of significant different compounds in *M. micrantha* under infection by *P. spegazzinii*.

**Supplementary Figure S2.**
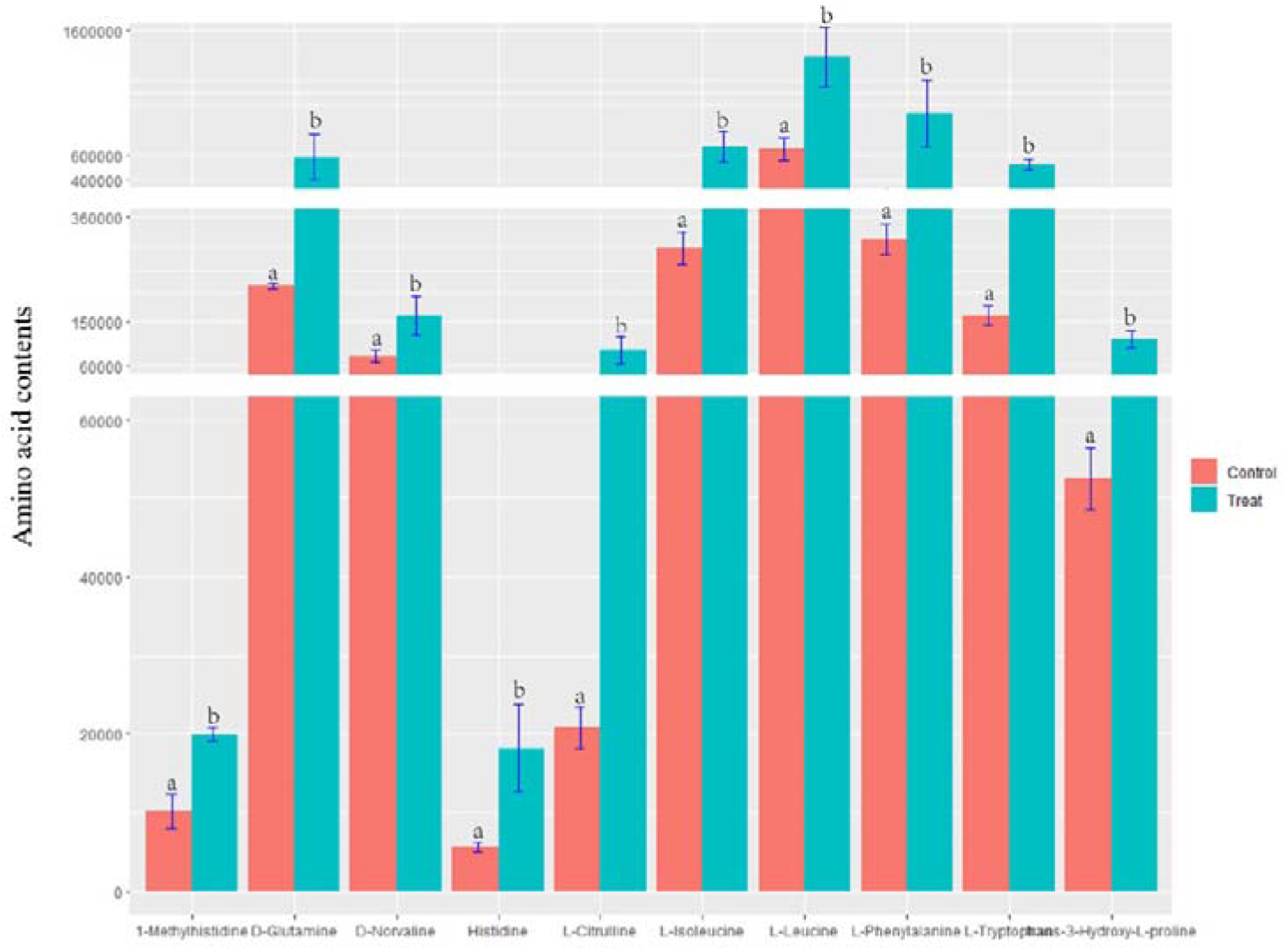
The difference of amino acid contents in *M. micrantha* leaves between the uninfected and infected by *P. spegazzinii*.

